# Foliar application of preparations as a method of protecting plants from the penetration of lead

**DOI:** 10.1101/2022.09.23.509154

**Authors:** M. Wierzbicka, K. Bodzon, A. Naziębło, Z. Tarnawska, M. Wróbel

## Abstract

Due to the contamination of soil with lead, there is still a danger of lead penetrating into our diet through crops. So far, no method has been developed to reduce the concentration of this toxic element in plants and to prevent it from entering the biological cycle. In this paper, an attempt was made for the first time to reduce lead concentrations in plants by using foliar calcium preparations.

This was based on the hypothesis that an increased amount of calcium in the plant would lead to the reduction in the amount of lead as the entry routes of calcium and lead are similar; therefore, an increase in the amount of calcium will result in the saturation of the routes through which lead enters cells (e.g. calcium channels). It should be clarified that the research was experimental. Three crop species (*Linum usitatissimum* L., *Solanum lycopersicum* L., *Cucumis sativus* L.) were tested at the organismal level, whereas the epidermis of *Allium cepa* L. was used to conduct tests at the cellular level. The InCa calcium transport activator (by Plant Impact) was selected for the test, followed by calcium nitrate. The preparations were administered foliarly. Lead, on the other hand, was applied to roots before adding lead nitrate into the mineral medium. The plants were cultivated hydroponically. The growth and development of seedlings, the concentration of lead in roots and shoots and the microscopic visualisation of lead in plant organisms and cells were studied. Ultimately, the InCa activator administered foliarly was shown to reduce the concentration of lead in plant organs by approximately 44%.

Further findings revealed that the mechanism of this process mainly resulted from the supply of calcium nitrate to plant leaves. A commercial calcium foliar fertiliser also showed a similar effect.

The potential to reduce the uptake of lead by crops by approximately 44% is a very satisfactory result. In addition, spraying plants with InCA biostimulant and calcium nitrate is environmentally friendly. This is cutting-edge research that was described for the first time in the present paper.

## 1. Introduction

Lead is a highly toxic heavy metal, found in nature mainly in the form of minerals, including galena. In the shallower soil layers, lead most often occurs as anthropogenic contamination. Despite restrictions on the use of this metal in production processes, significant amounts of this element are still accumulated in soils all over the world. Moreover, as a result of human activities, each year hundreds of tonnes of lead enter the environment mostly from the mining and smelting industries (Lis and Pasieczna, 1995; Cegiełkowska et al., 2015).

We have been conducting research on lead management in various plant species for many years. (Wierzbicka, 1998; Wierzbicka, 1999b; Wierzbicka, 2002; Baranowska-Morek and Wierzbicka, 2004). Routes of lead uptake (Wierzbicka, 1987a; Wierzbicka, 1987b; Wierzbicka, 1995), accumulation of lead in tissues, cell walls and vacuoles (Chow et al., 1970) and mechanisms of its detoxification (Wierzbicka, 1994; Wierzbicka, 1995; Wierzbicka, 1998). These processes ultimately lead to the safe accumulation of lead in plant tissues, which is dangerous for animals and humans that consume them (Wierzbicka and Antosiewicz, 1993). For this reason, it is of the utmost importance to limit the accumulation of Pb^2+^ ions in crops intended for animal feed and direct human consumption. So far, researchers have not succeeded in developing effective methods to reduce the uptake of lead from the soil by plants. This is a serious threat as good methods for eliminating heavy metals from soils still have not been worked out. The available physical and chemical techniques require serious interference with the soil ecosystem, e.g., by pouring chelating compounds on the soil or removing the entire topsoil (Hettiarachchi and Pierzyński, 2004) and roasting the soil at high temperatures. As a result, these methods are not satisfactory. It may be possible to reduce the bioavailability of heavy metals to plants by liming the soil, but this process significantly increases soil pH, which is detrimental to the growth and development of most crops (Tlustos et al., 2006; He et al., 2021).

The aim of the research conducted was to develop a method that would reduce the penetration of lead into the body of a plant. Such an attempt was made on the basis of available test results suggesting that calcium ions can effectively inhibit the absorption of heavy metals by the body (Skórzyńska-Polit, 1998; Lock et al., 2007; Tian et al., 2007; Wang and Song, 2009). This fact is explained by the ability of Ca^2+^ ions to displace toxic elements from various compounds, relieve metal-induced oxidative stress and stimulate the photosynthetic activity of plants. The plant species tested showed an increase in metal resistance along with an increase in calcium concentrations, and some even developed a higher tolerance to Pb^2+^ ions (Lane et al., 1978; Garland and Wilkins, 1981; Chadzinikolau et al., 2011).

The existence of a relationship between calcium and lead ions is also confirmed by studies conducted on animal organisms; lowering the calcium content in the diet of test rodents and birds leads to an increased absorption of lead ions, whereas the supplementation of calcium ions stimulates the antioxidant activity and mitigates the toxic effects of lead (Six and Goyer, 1970; Dauwe et al., 2006; Jaya Prasanthi et al., 2010).

We paid particular attention to the InCa preparation. It is a foliar biostimulant produced by Plant Impact. The manufacturer considers it to be a calcium transport activator. The action of the InCa activator is based on supplying the plant with calcium in the form of an easily soluble salt (nitrate(V)) and stimulating its transport to those parts of the plant that are most deficient in this element. This requires the stimulation of the proton pump that enables calcium ions, which move in plant organisms mainly via the apoplastic pathway, to be directed into cells. For this purpose, molecules of auxin, i.e. hormone responsible for activating ATPases in the cell membrane and increasing its permeability to Ca^2+^ ions, were included in the fertiliser formulation (López et al., 2005). To maintain hormonal balance, it also contains cytokinin (diphenylurea). The hormonal components are extracted from algae, as is the case with many other foliar calcium or magnesium fertilisers. Not only do such extracts facilitate the penetration of leaf tissue, but they also contain numerous poly- and oligosaccharides, vitamins, and micronutrients that contribute to the resistance of cultivated plants (Gawrońska, 2008).

The aim of the research conducted was to look for opportunities to reduce the uptake of lead by crops.

This paper discusses the following hypothesis: Since lead enters plants via a similar route to that of calcium, it was assumed that increasing the amount of calcium would reduce the uptake of lead by plants.

In conducting this research, particular attention was drawn to modern agricultural preparations, namely biostimulants, which are used to improve crops and storage of fruit and vegetables. Such products stimulate the natural physiological processes of plants and thus ensure their better growth and development. They are often calcium transport activators.

## 2. Material and Methods

### 2.1. Tests at the organismal level

Tests at the organismal level were carried out on plants at the seedling stage. The following species were studied: *Linum usitatissimum* L., *Solanum lycopersicum* L., *Cucumis sativus* L.

### 2.2. Plants at the seedling stage

The plants at the seedling stage were cultivated hydroponically. For this purpose, 20 seeds of each plant were sown on filter paper with 5 ml of distilled water at 23ºC. After germination, seedlings were transferred to culture containers (with a capacity of 5 litres). They were attached to glass racks in such a way that the roots were immersed in a mineral medium - Knop’s medium (Knop, 1865). Lead nitrate Pb (NO3)2 was added to some containers, usually at a concentration of 3 mg/l of lead. In this way, lead was administered to the roots of the plants, whereas the calcium transport activator, including InCa, was applied foliarly by spraying the plants at a concentration of 0.01% to 0.006% every 2-3 days. Control plants were those cultivated in the mineral medium without additional preparations.

The incubation of the plants lasted from 5 to 14 days. The plants were grown in a growing room with a photoperiod of 16h during the day and 8h at night, at 23ºC during the day and at night, with 12,000-140,000 lux of light. As many as 50 plants were grown in one experimental combination, 300 plants in one experiment, and a total of 10,000 plants were examined.

### 2.3. Biometric observations

Morphological observations of the plants were carried out during all tests conducted at the organismal level. The length of the main root (Wilkins, 1978) and the shoot, as well as the size of the first leaf, were measured on specific days of hydroponic culture.

At the end of the experiments, the plants were divided into underground and aboveground parts. The roots were washed in distilled water using an ultrasonic cleaner and their fresh and dry weight were determined. Then, they were subjected to chemical analysis.

### 2.4. Chemical analysis

The plant samples prepared in this way were subjected to chemical analyses that were carried out using the FAAS method (Bulska and Krata, 2006). Lead and calcium concentrations were determined in individual parts of the plants. The samples were mineralised using the Milestone Ethos microwave mineraliser in 69% HNO3, analytical grade, with an addition of 35% H2O2, analytical grade. The parameters were determined using the Sollar M6 equipment. Some of the analyses were carried out using the flame technique with background correlation by means of a deuterium lamp. The second part of the analyses was performed using the graphite cuvette technique with Zeeman background correction.

The reference material was Virginia Tabacco Leaves CTA-VTL-2.

### 2.5. Visualisation of lead in tissues and cells

#### 2.5.1. Tissue visualisation of lead

The dithizone method was used to visualise lead in plant tissues. For this purpose, plant cells were placed in a 0.05% acetone solution of dithizone (diphenylthiocarbazone) (Abratowska, 2013). The staining time was 24 hours. The tissues were then observed under the light microscope. A change in the colour of the tissues to red was indicative of the presence of lead. At the tissue level, lead was visualised based on cross sections through the root and shoot *C. sativus*, using a scanning electron microscope with X-ray microanalysis. Sections through the plants were dried and then observed under the scanning electron microscope (Phenom ProX) with X-ray microanalysis. Area microanalysis yielded a map of lead distribution in plant tissues.

#### 2.5.2. Ultrastructural visualisation of lead

Ultrastructural observations were carried out using electron microscopy techniques. Root tips were taken from the relevant experimental variants. They were fixed in 2.5% glutaraldehyde in cacodylate buffer for 2h and then in 1% osmium tetroxide for 16 hours. After dehydration, the material was saturated with Epon-Spurr epoxy resin (Antosiewicz and Wierzbicka, 1999).

Ultra-thin sections were cut using an ultramicrotome (Power Tome XL) with a diamond knife. The ultrastructure of the cells was observed under the transmission electron microscope.

#### 2.5.3. Tests at the cellular level

Tests evaluating the effect of the InCa activator on the toxicity of lead were conducted at the cellular level. The experiments were carried out on epidermal cells of storage leaves of the *Allium cepa* L. (onion) scale. The epidermis was remove with tweezers from the concave side of the scale. Each fragment of the epidermis was divided into two parts. The first part was put in a solution of the tested biostimulants (Ametros, Voyager, BioCal, and InCa) (Table 1) for 15 minutes and then transferred to a lead nitrate solution for 3 hours and 45 minutes. The other part of the epidermis was placed in the respective biostimulants for 15 minutes and then in water for 3 hours and 45 minutes. After the incubation period, the epidermis was washed in distilled water. Microscope slides were prepared and observed under the light microscope, fluorescence microscope, and confocal microscope.

**Tab 1.**
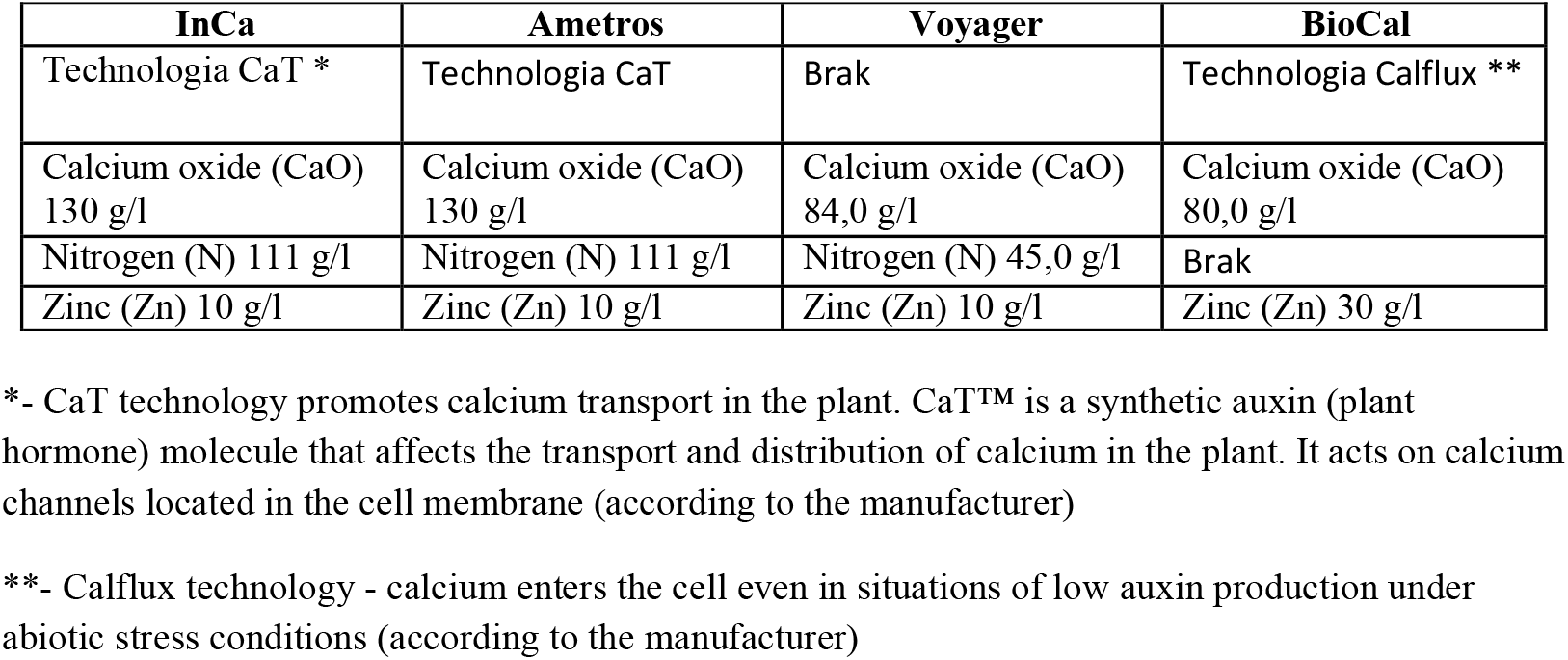
Comparison of the composition of the tested biostimulants by manufacturer: InCa, Ametros- by Plant Impact Plc, BioCal- by BioArgis, Voyager- by Engage Argo Europe Company

#### 2.5.4. Measurement of the metabolic activity of cells

To determine the toxic effects of lead on cells, the movement of the cell cytoplasm, which reflects the metabolic activity of cells, was assessed. After the period of incubation in the respective biostimulants and then in lead nitrate, microscope slides were prepared and observed under the phase-contrast light microscope (Nikon OPTIPHT-2). They were used to count cells with the visible movement of the cytoplasm. A total of 48 cells were observed on one slide (8 fields of vision each). Twenty-four slides were analysed in each experimental combination, i.e. a total of 1,152 cells. All experiments were repeated twice.

#### 2.5.5. Measurement of the damage to the cell membrane

The number of living and dead cells is indicative of the degree of damage to plasmodesmata (Przedpelska, 2004 after Broda, 1971). After the incubation of cells in the biostimulants and in the lead nitrate solution, the epidermis was immersed in a neutral red solution for approximately 5 minutes. Observations were carried out in the same way as those of the movements of the cytoplasm. All experiments were repeated twice.

#### 2.5.6. Visualisation of lead in cells

To visualise lead in the cells, Leadmiu™ Green AM dye fluorescent probe was used. It can also be used on living cells. The probe is specific for the detection of lead and cadmium in cells (Molecular Probes). The excitation and emission maxima for Leadmiu™ Green are 490nm and 520nm, respectively (Molecular Probes). Staining was carried out according to the method (Jiang et al., 2014). Observations were performed using the NIKON A1R MP confocal microscope equipped with VIS LASERS: 404 nm, 488 nm, 561 nm, and 638 nm, a 32-channel spectral detector with a resolution of 2,5,6 or 10 nm (Virtual Filter mode), and a CCD DS-5Mc camera with a resolution of 5Mp at the excitation of 488 nm and with a barrier filter of 590/50 nm (Jiang et al., 2014).

#### 2.5.7. Statistical analysis

The results obtained during the tests were presented as an arithmetic average with standard deviation. The graphical representation was prepared in Microsoft Excel. The statistical analysis was carried out using the STATISTICA software, version 13.1. (STATSOFT, INC.). The non-parametric Kruskal-Wallis test, for multiple independent samples significance level α =0.05, was used to compare several groups of the variants tested.

## 3. RESULTS

### 3.1. Selection of biostimulants for testing

As there is a number of biostimulants on the market that affect the metabolism of calcium in plants, their effect was assessed using the epidermal test of *Allium cepa* L. (Przedpelska, 2004). The tests were carried out using commercial products that are calcium transport activators, such as InCa, Ametros, Voyager, and Biocal (Arysta Life Science, 2014; Plant Impact, 2018; BioArgis brochure, 2020). The results are presented in Fig. 1 and Fig. 2. All biostimulants tested showed positive effects, i.e., a reduction in the toxic effects of lead on cells. There was an increase in metabolic processes, as indicated by an increased movement of the cytoplasm in cells (Fig. 1), and a decrease in damage to plasmodesmata as shown by an increased number of living cells (Fig. 2). Taking the degree of lead toxicity reduction into account, the biostimulants can be listed in the following order: InCa (by 57%), BioCal (by 56%), Ametros (by 47%), Voyager (by 40%). Although all biostimulants tested showed a reduction in the toxicity of lead, InCa was the most effective one.

**Fig. 1.**
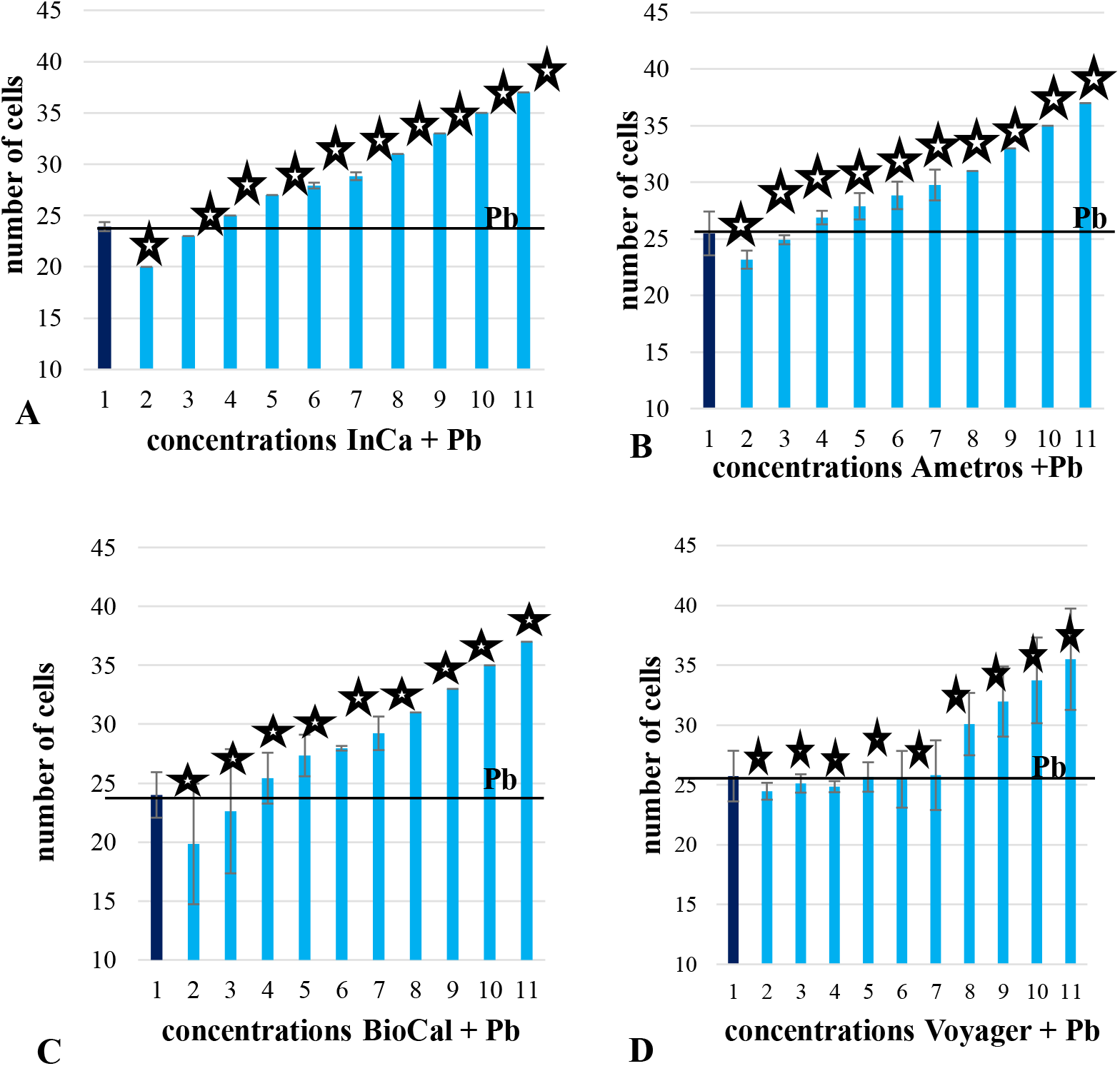
Cytoplasmic movements in *Allium cepa* L. epidermis cells incubated in 20 mg/l Pb (1) and in combination with biostimulants with the following concentrations: (2) – 0.2 mg/l; (3) – 0.02 mg/l; (4) – 0.01 mg/l; (5) – 0.001 mg/l; (6) – 0.0001 mg/l; (7) – 0.0005 mg/l; (8) – 0.00005 mg/l; (9) – 0.00001 mg/l; (10) – 0.000005 mg/l; (11) – 0.000001 mg/l. Asterisks indicate statistically significant differences with respect to lead. The applied biostimulants was A – InCa, B – Ametros, C – BioCal, D – Voyager.

**Fig. 2.**
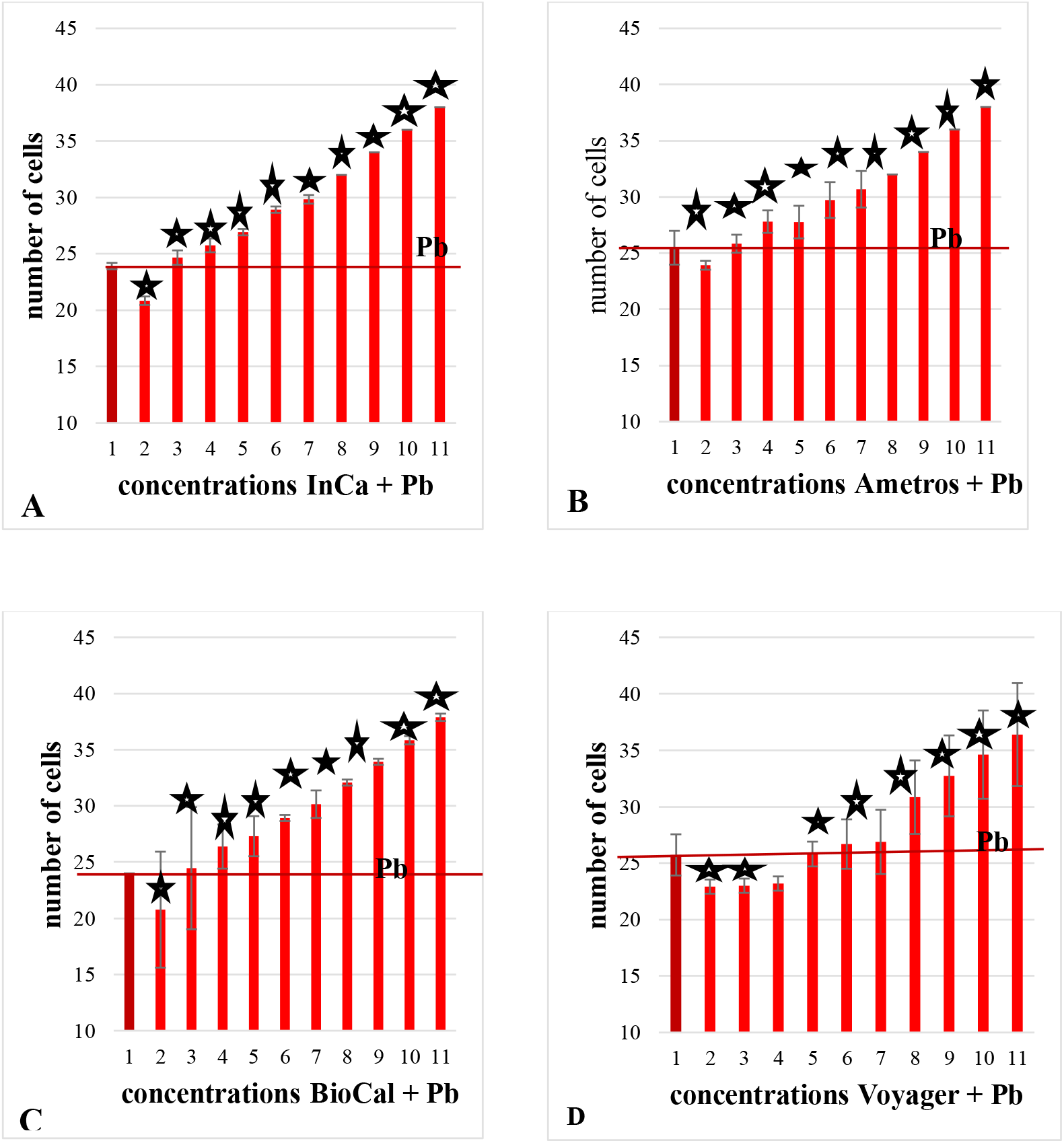
Number of living cells (mean of 1,152 cells) among 48 *Allium cepa* L. epidermis cells incubated in 20 mg/l Pb and in combination with biostimulants with the following concentrations: (2) – 0.2 mg/l; (3) – 0.02 mg/l; (4) – 0.01 mg/l; (5) – 0.001 mg/l; (6) – 0.0001 mg/l; (7) – 0.0005 mg/l; (8) – 0.00005 mg/l; (9) – 0.00001 mg/l; (10) – 0.000005 mg/l; (11) – 0.000001 mg/l. Asterisks indicate statistically significant differences with respect to lead (1). The applied biostimulants was A – InCa, B – Ametros, C – BioCal, D – Voyager.

### 3.2. Selection of metal for testing

The next stage of the research involved microscopic observations of several metal compounds to see which compound would reduce its toxicity towards *Allium cepa* cells under the influence of the InCa biostimulant.

The tests included the compounds of four metals in the form of nitrate salts, three of which are divalent: Pb, Zn, Cd, and one is monovalent: Tl. The results are presented in Fig. 3. In this respect, the test results clearly show that the InCa biostimulant reduced the toxic effects of lead by 48,5%, cadmium by 20%, zinc by 22%, and thallium by 0%.

**Fig. 3.**
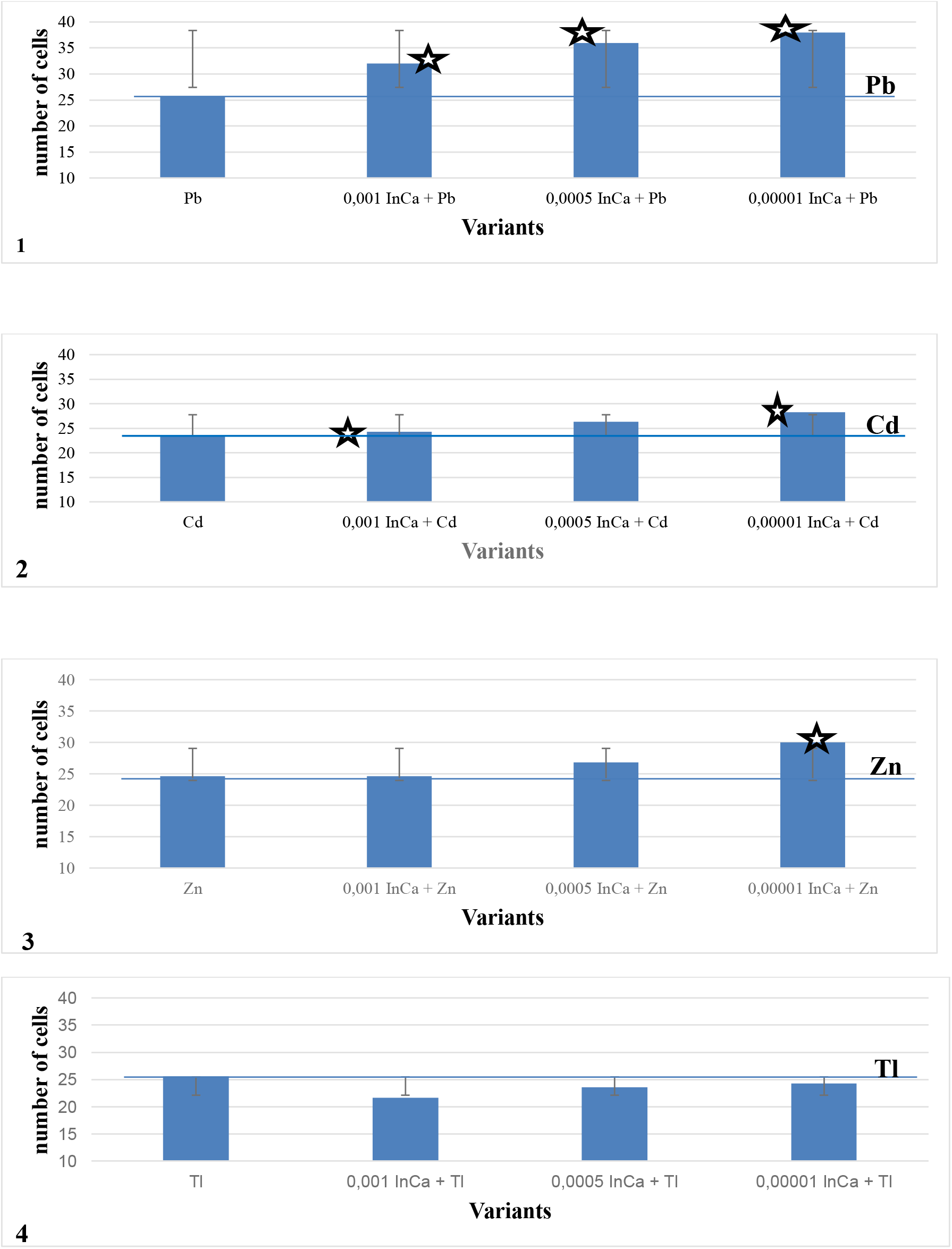
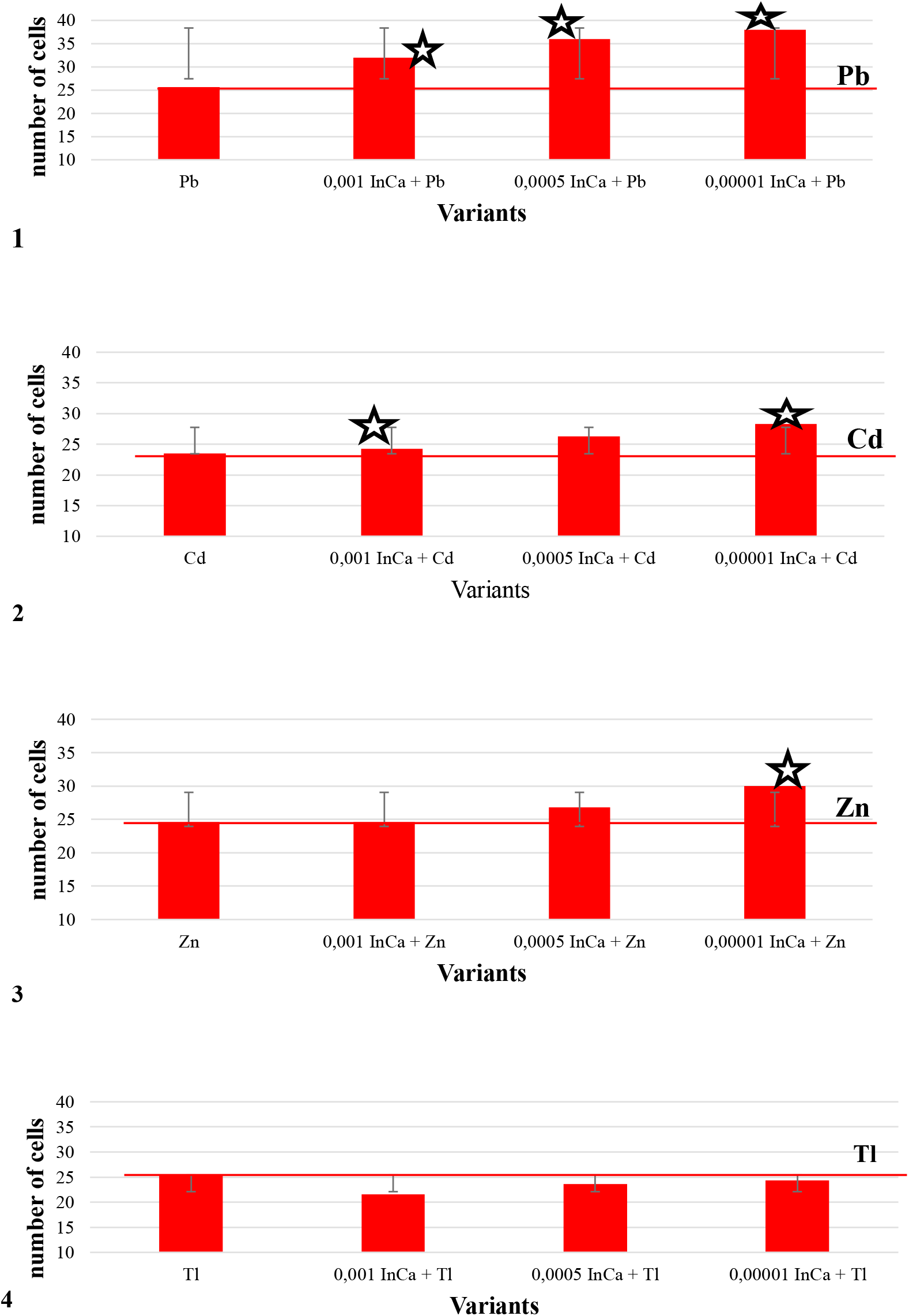
Comparison of the reduction in toxic effects of metals: 1 – Pb (20 mg/l); 2 – Cd (5 mg/l); 3 – Zn (5 mg/l); 4 – Tl (1 mg/l) after application of the InCa biostimulant. The test of *Allium cepa* L. epidermis was performed. Three concentrations of biostimulant were used. Metals were administered as nitrates. a – number of cells with cytoplasmic movements b – number of living cells. Asterisks indicate statistically significant differences with respect to metals.

Due to the fact that the strongest effect was observed for lead, the authors decided to choose this element for further testing.

### 3.3. Selection of plant species for testing

Four crop species, differing in calcium status, were selected for testing (Antosiewicz, 1993). These were: *S. lycopersicum* L., *M. sativa* L., *C. sativus* L., *L. usitatissimum* L.

The above plant species were grown in a medium with a reduced content of calcium. Based on the results obtained, *S. lycopersicum* L. tolerates calcium deficits well, *C. sativus* L. and *L. usitatissimum* L. show medium tolerance to calcium deficits and *M.sativa* L. is sensitive to calcium deficits. These results are consistent with the data obtained by Antosiewicz (1993). The tolerance of the plant species tested to lead in combination with a reduced amount of calcium in the medium was also investigated. The results are presented in Fig. 4.

**Fig. 4.**
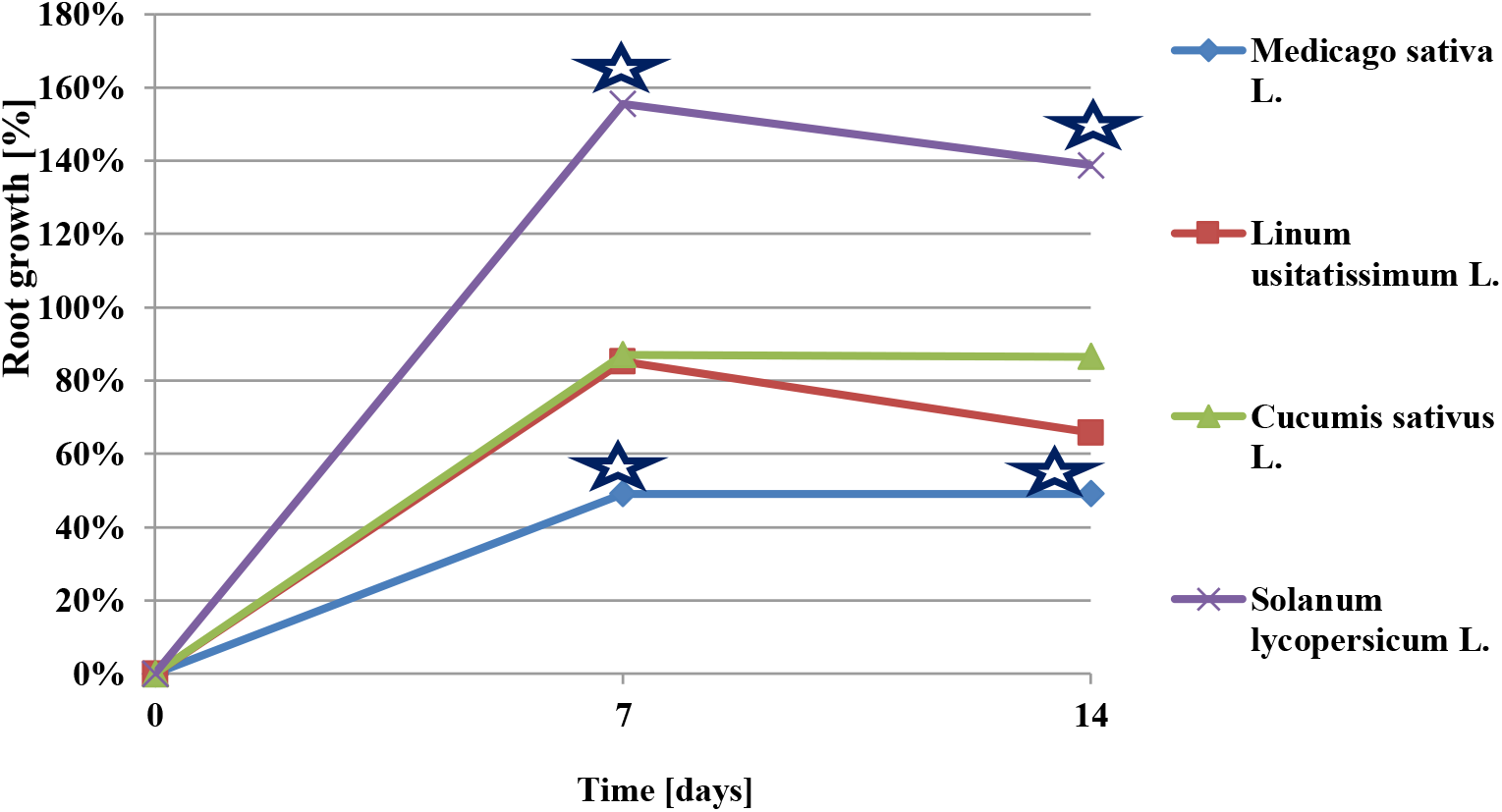
Root growth of plants: *Medicago sativa* L; *Linum usitatissimum* L.; *Cucumis sativus* L.; *Solanum lycopersicum* L., which were grown in a medium with reduced calcium content (1% Ca) and with added lead at 20 mg/l. Asterisks indicate statistically significant differences with respect to lead.

Based on the above-mentioned findings, *S. lycopersicum* L. and *Cucumis sativus* L. were selected for further testing as species demonstrating a high and medium tolerance to lead and high and medium tolerance to calcium deficits (respectively).

### 3.4. Effects of the InCa biostimulant

The InCa biostimulant is a type of foliar fertiliser. Several-day-old seedlings of the plants tested were sprayed with the InCa biostimulant. Their response to this factor is shown in Fig. 5. The dose of InCa (0.01%) was adjusted so that it did not inhibit the growth of the plants and was well tolerated by them, whereas the dose of lead applied to the roots of the plants tested was set at 3 mg/L. This was the lowest possible dose that did not inhibit the growth of the plants, making it the closest to natural conditions (Fig. 5). The dose was well tolerated by the plants.

**Fig. 5.**
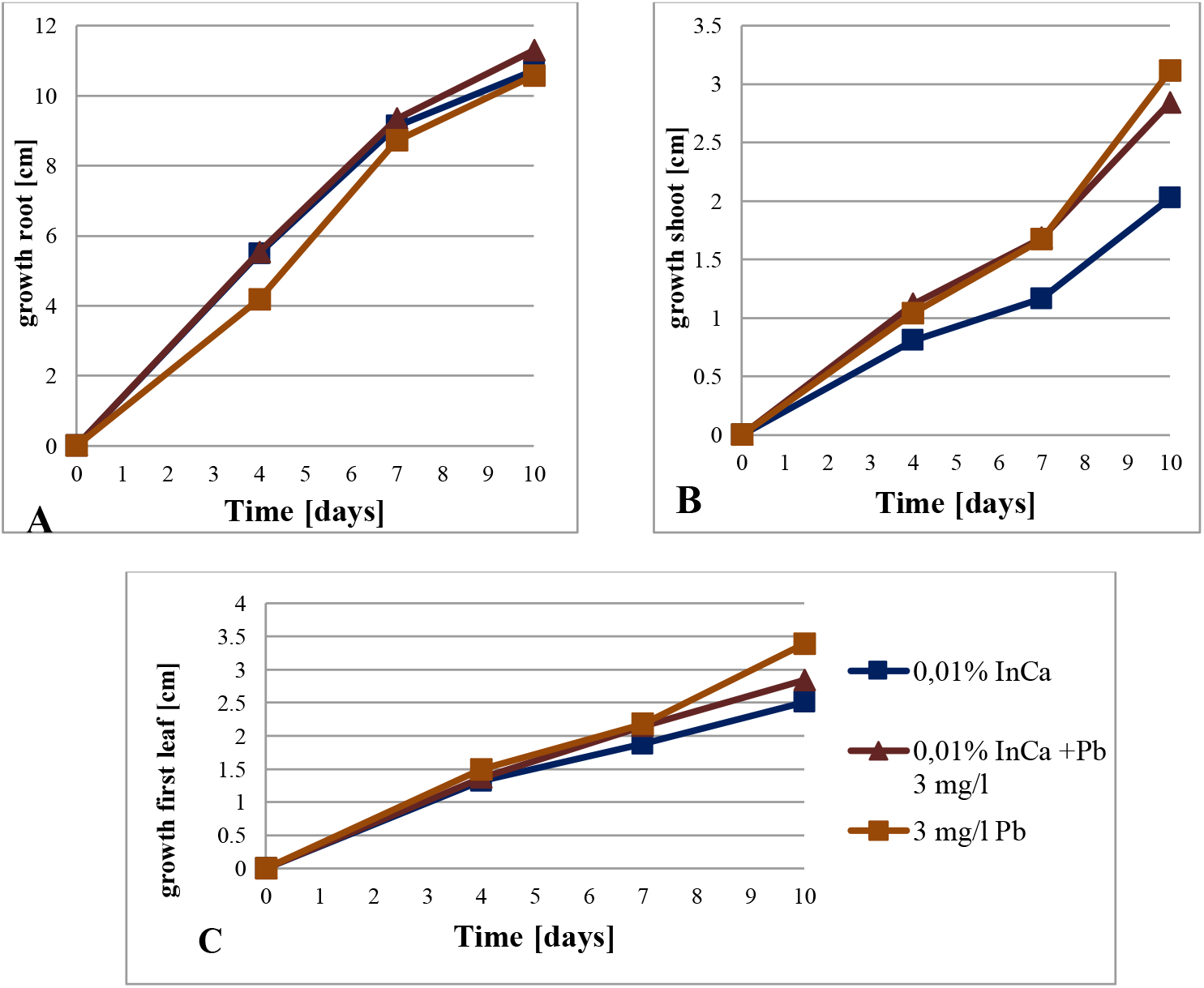
Growth of *Cucumis sativus* L. plant organs: A – root, B – shoot, C – first leaf. Plants were treated with lead administered to the roots at 3 mg/l and treated foliarly with the activator InCa at 0.01%. Both lead and InCa have no inhibitory effect on plant growth. No statistically significant differences.

### 3.5. Lead concentration in plants after the foliar application of the InCa biostimulant

Tests were carried out to determine the concentration of lead in plants after the foliar application of the InCa biostimulant and with a simultaneous administration of lead to the roots. After 24 and 48 hours application of the InCa, there was a reduction in lead concentrations for both test species (Fig. 6 A, B). And, thus, the concentration of lead in the roots of *S. lycopersicum* L. was reduced by 47% and by 20% and *C. sativus* L. by 33% and by10.8%. The concentration of lead in the aboveground parts of the plants, after the foliar application of InCa, in *S. lycopersicum* L. was reduced by 36% and by 49% (Fig. 6 A, B).

**Fig. 6.**
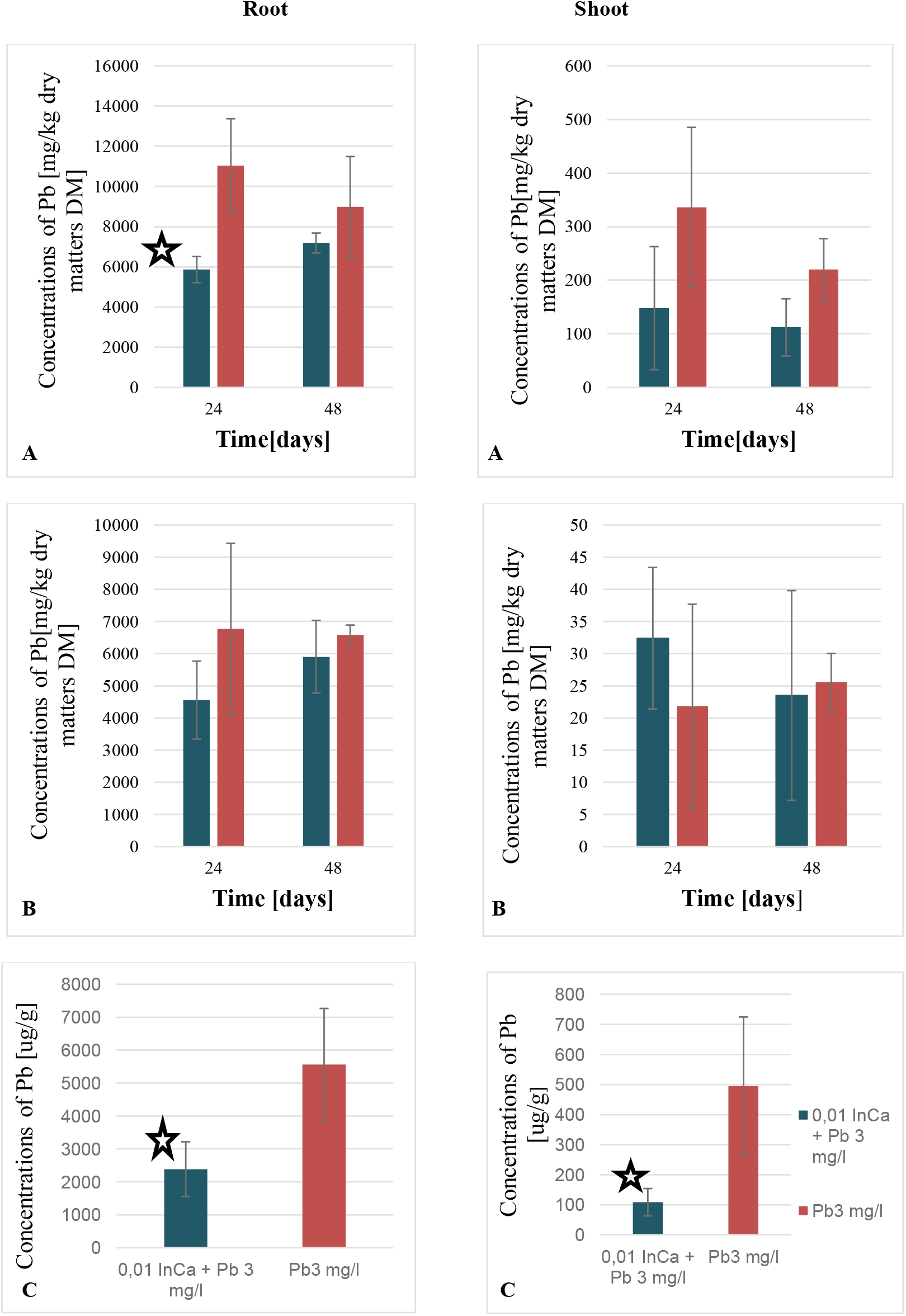
Comparison of lead concentrations in roots and shoots of plants treated with lead with plants treated with InCa+Pb (A – *Solanum lycopersicum* L.; B – *Cucumis sativus* L.; C – *Linum usitatissimum* L. after 10 days) treated with lead administered to the roots (3 mg/l) and after foliar administration of InCa at 0.01% after 24 and 48 hours. Asterisks indicate statistically significant differences with respect to lead.

Analogous observations were made for the third species studied. After 10 days of application of InCa in *L. usitatissimum* L. (Fig. 6 C), there was a 57% reduction in root and 78% reduction in shoot lead concentrations.

In conclusion, the foliar application of the InCa activator resulted in a reduction of lead concentrations in roots by an average of 33.4% and in shoots by 54.3%, i.e. approx. 44%.

### 3.6. Visualisation of lead in plant parts and cells

The question arises about the distribution of lead in plant tissues and cells after the application of the InCa activator. There are several possible answers to this question. Fig. 7 presents the visualisation of lead in *C. sativus* L. roots treated with InCa, which was applied foliarly, compared to the control (Fig. 7A). Lead was taken up by the entire root surface (Fig. 7B). The staining of the roots of plants treated with lead and InCa applied foliarly (Fig. 7C-arrow) was weaker compared to plants treated with lead alone (Fig. 7B).

**Fig. 7.**
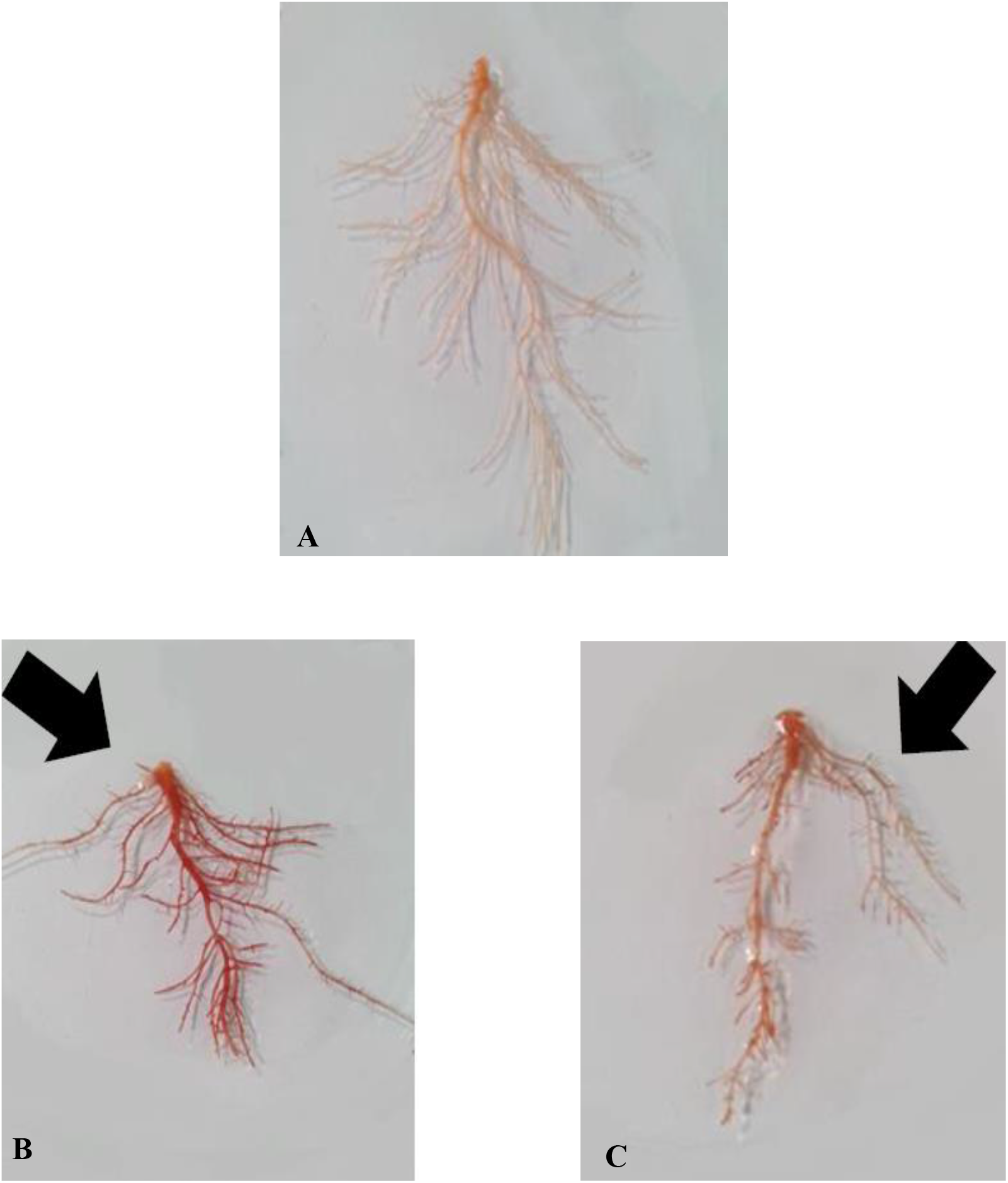
Visualisation of lead in *C. sativus* L. roots stained with dithizone solution after 4 days of plant treatment. A – control plants; B – plants treated with 6 mg/l Pb; C – plants treated with 0.2% InCa and 6 mg/l Pb. Red colouration indicates the presence of lead. (Kraskiewicz, 2016)

The ultrastructural localisation of lead in *C. sativus* L. root cells revealed its presence in cell walls (Fig. 8A). In contrast, after the foliar treatment of the plants with InCa (application of lead to the roots), the lead deposits were fewer but large in size (Fig. 8B).

**Fig. 8.**
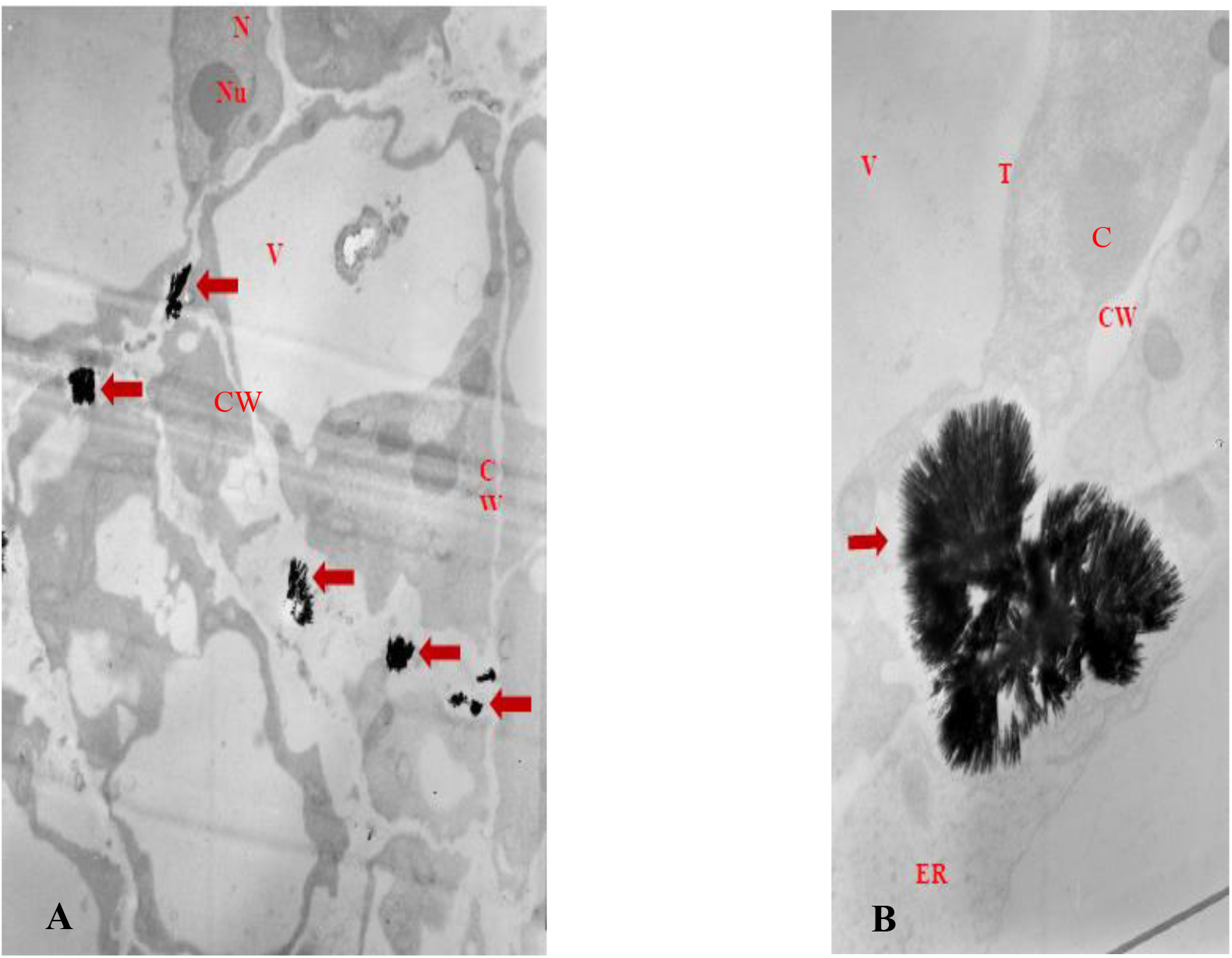
Ultrastructural visualisation of lead in meristematic cells of *C. sativus L*. root by transmission electron microscopy: A – cells after 4 days of plant treatment with 6 mg/l Pb, above 3400x; B – cell after 4 days of plant treatment with 0.2% InCa and 6 mg/l Pb, above 8000x. Red arrows show lead deposits. CW – cell wall, V – vacuole, N – nucleus, Nu – nucleolus, C – cytoplasm. (Kraskiewicz, 2016)

The localisation of lead was also carried out at the cellular level using the fluorescent probe (Fig. 9A) with respect to the *A. cepa* L. epidermis. The research revealed that lead was mainly accumulated in the cytoplasm of the cells as shown by the green fluorescence (Fig. 9B). However, after the additional application of InCa, less lead entered the cells. This is indicated by the faint staining of the cells’ cytoplasm (Fig. 9C). The control is shown in Fig. 9A.

**Fig. 9.**
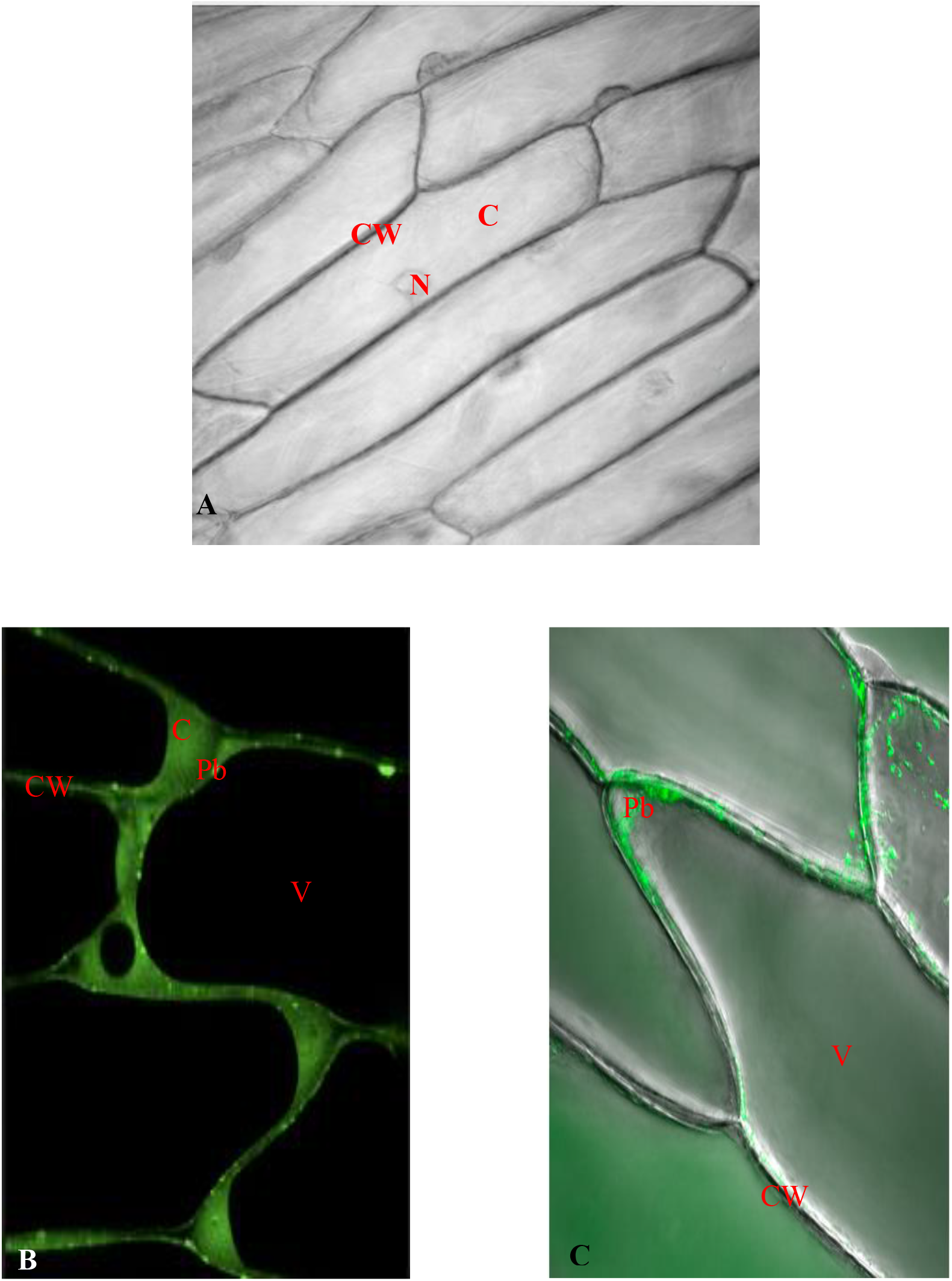
*A. cepa L*. epidermal cell. Lead in the cells was visualised with the fluorescence probe Leadium™ Green. A – control; B – 100 mg/l Pb; C - 0.01% InCa + 100 mg/l Pb. Green fluorescence indicates the presence of lead. CW – cell wall, V – vacuole, N – nucleus, C – cytoplasm

The above-mentioned research indicates a reduction in lead uptake by *C. sativus L*. plants, both at the whole organ level and the cellular level (cytoplasm and epidermis), as well as cell ultrastructure (root cell wall). A reduction in lead content after InCa treatment was demonstrated in all compartments – i.e., both in the apoplast (cell wall. Fig. 8) and symplast (cytoplasm, Fig. 9).

### 3.7. Which component of the InCa activator is responsible for reducing the uptake of lead by plants?

The InCa activator was developed by the manufacturer to improve the quality of crop yields. The current purpose of its use in agriculture is different from that sought in this study. This research leads to a reduction in lead uptake by plants. The question remains as to the mechanism of this process.

For this purpose, the individual components of the InCa biostimulant were tested to see which one was responsible for reducing lead uptake. In this way, calcium nitrate, DPU and IAA hormones, citric acid and various combinations of these compounds were tested. Finally, it was found that the lead-reducing properties in *C. sativus L*. roots were demonstrated by calcium nitrate. The results are presented in Fig. 10. After the application of calcium nitrate the reduction in lead concentrations was 73% however, after InCa was 67%.

**Fig. 10.**
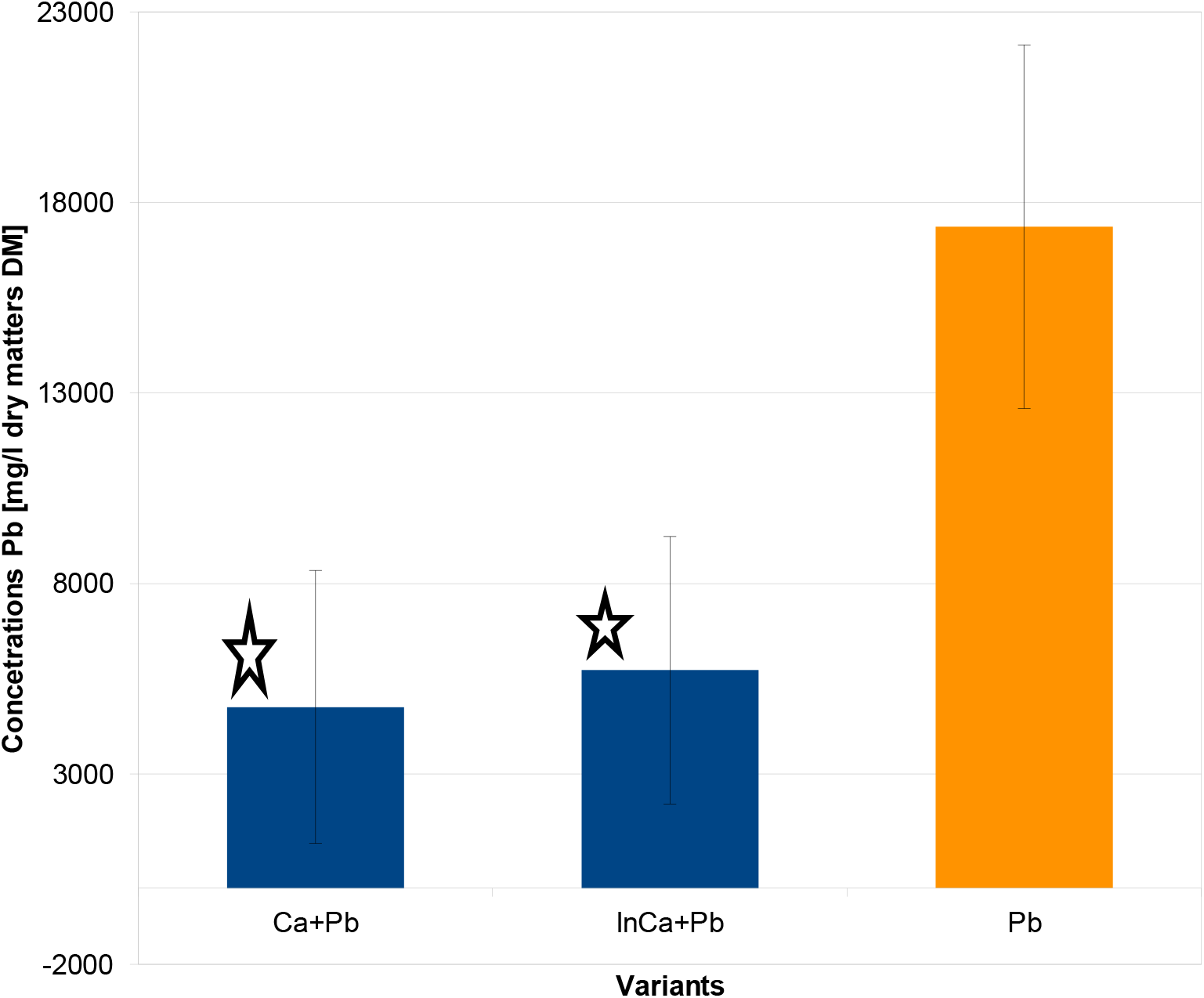
Lead concentrations in roots of *C. sativus* L. seedlings treated foliarly with calcium nitrate solutions or InCa and treated with lead administered to the roots. Asterisks indicate statistically significant differences from plants that were not treated foliarly with the tested preparations.

Visualisation of lead in *C. sativus* L. roots and shoots treated with lead administered to the roots and treated foliarly with calcium nitrate was performed on cross-sections by scanning electron microscopy. Elemental distribution was performed using X-ray microanalysis. Fig. 11 shows a map of lead distribution on the cross-section through the vascular bundles. In contrast, Fig. 12 shows the difference in terms of lead concentrations between plants treated with lead and calcium nitrate + lead. The analysis provided an estimate of lead concentrations in tissues (Fig. 13). In this way, foliar application of calcium was found to reduce lead accumulation in the vascular bundles by 89% in the root parenchyma by 91% in the epidermis by 78% -by an average of 86% (Fig. 13).

**Fig. 11.**
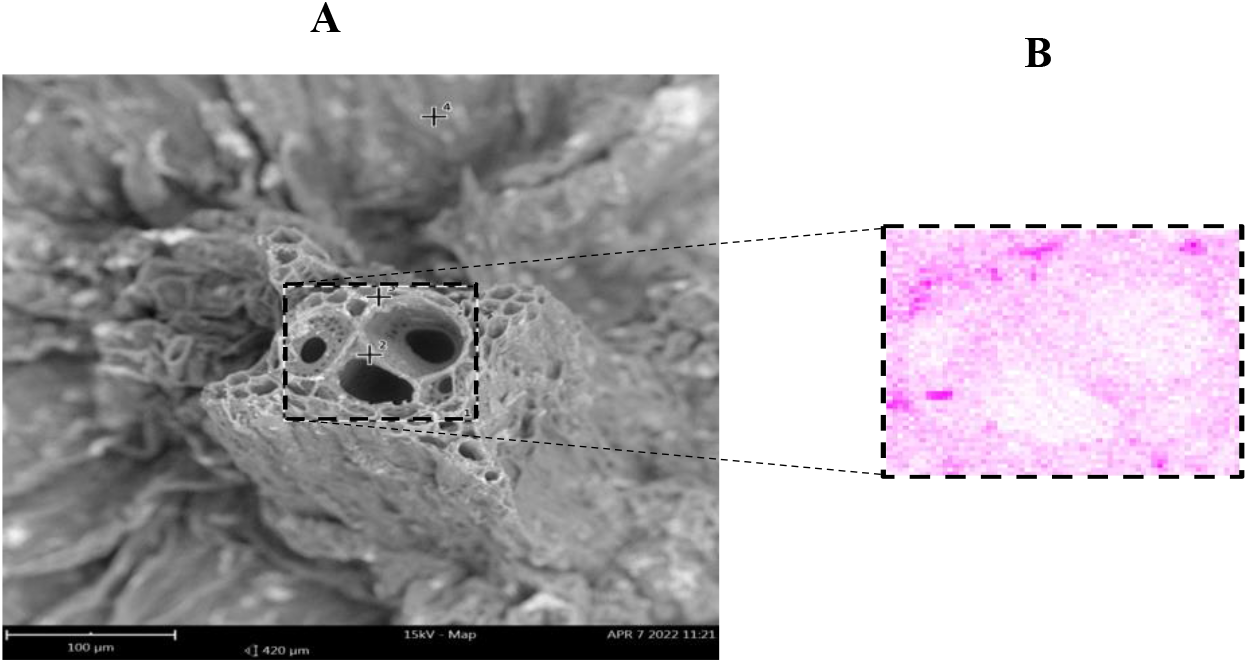
Distribution of lead in the vascular bundle of roots of *C. sativus* L. seedlings treated with 10 mg/l Pb. A – cross-section through the root; B – map of lead distribution. Application of an area X-ray microanalysis in the vascular bundle. Scanning electron microscope image with X-ray microanalysis performed. Red colour indicates the presence of lead.

**Fig. 12.**
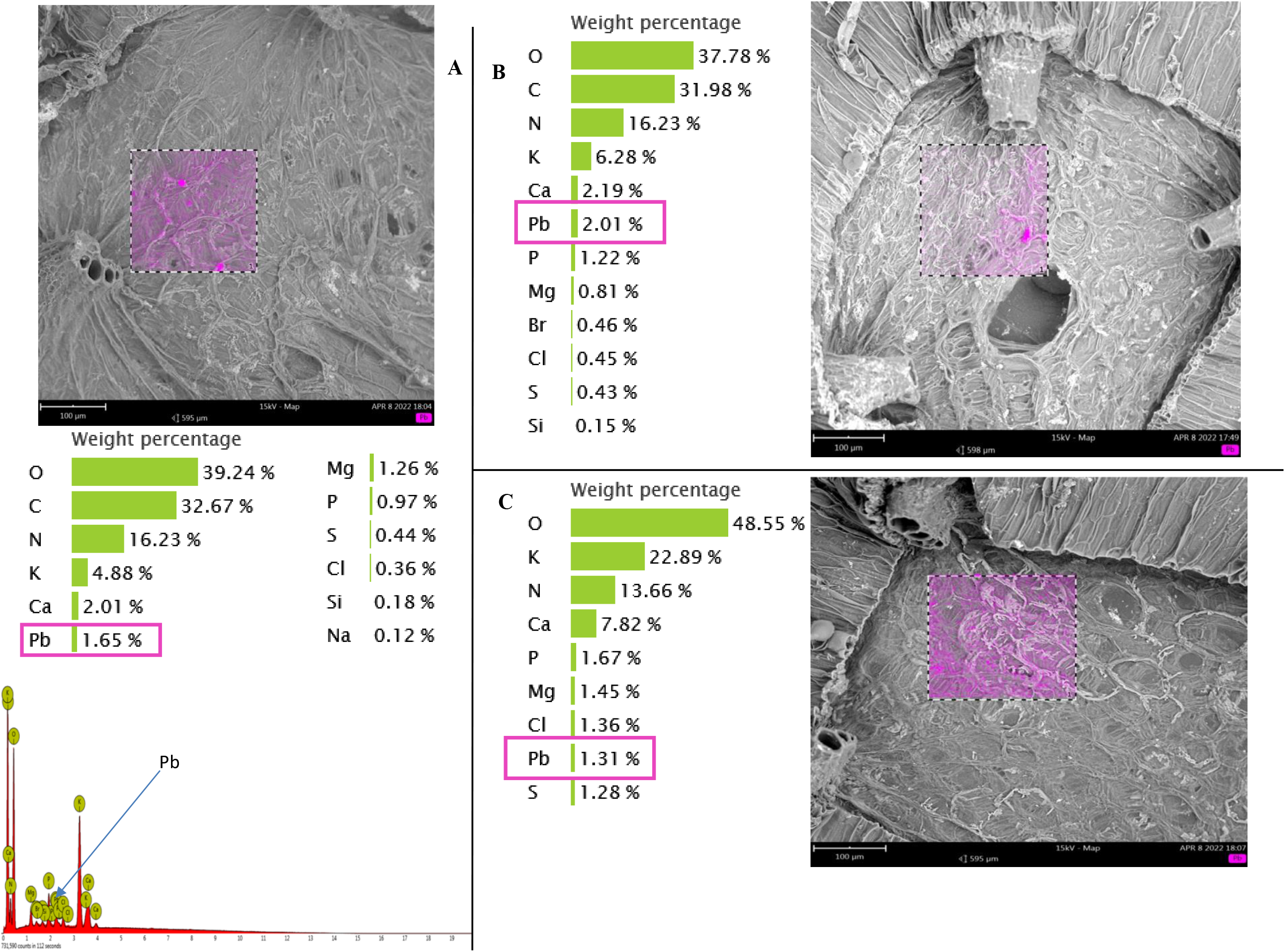
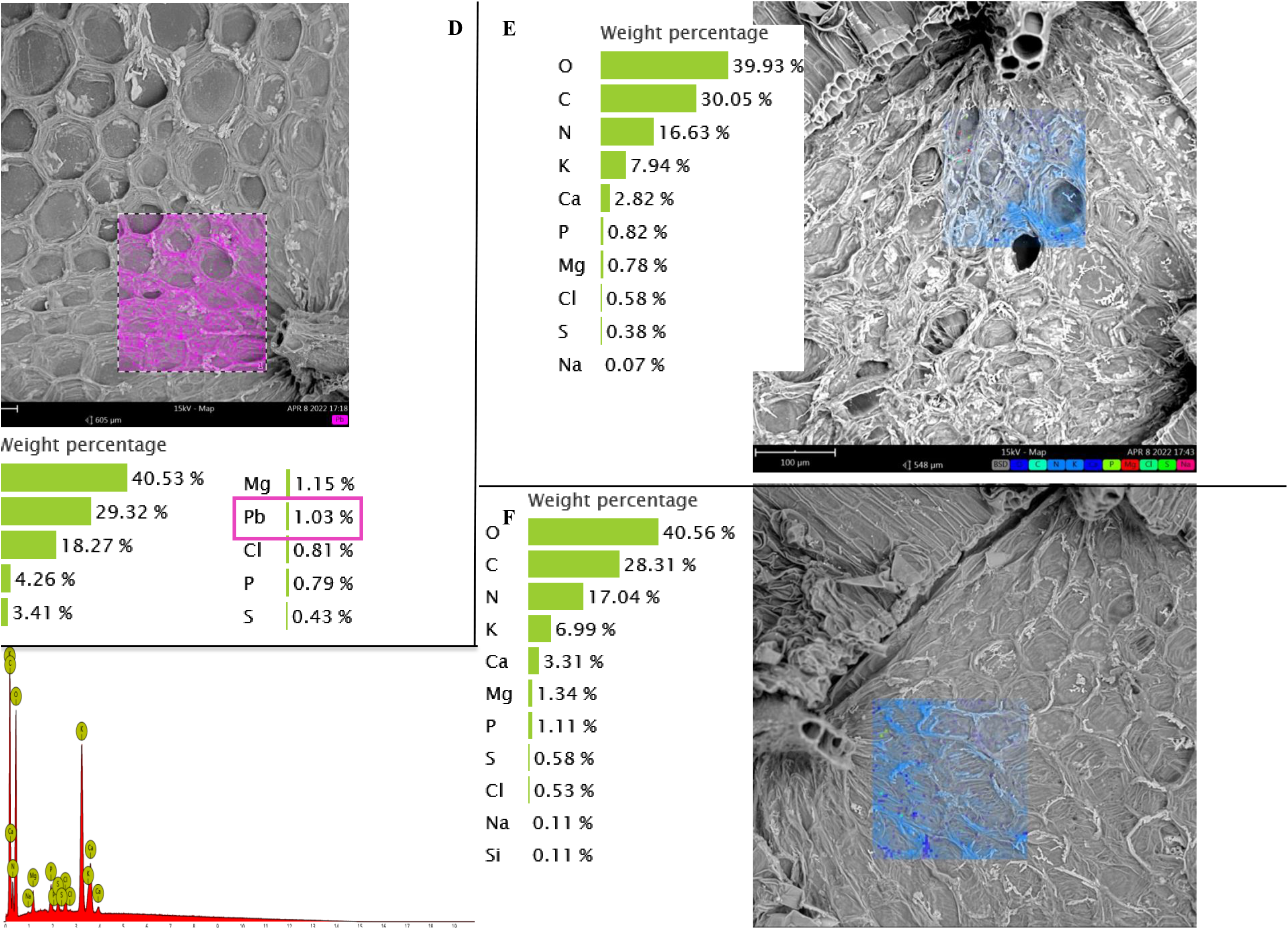
Distribution of lead in *C. sativus* L. shoots on cross-sections. Area X-ray microanalysis. A – plants treated with lead administered to the roots; B – plants treated with lead administered to the roots and treated foliarly with calcium nitrate. Red shows the distribution of lead and blue shows no lead detected. Figures and tables show the elemental analysis of the area under study.

**Fig. 13.**
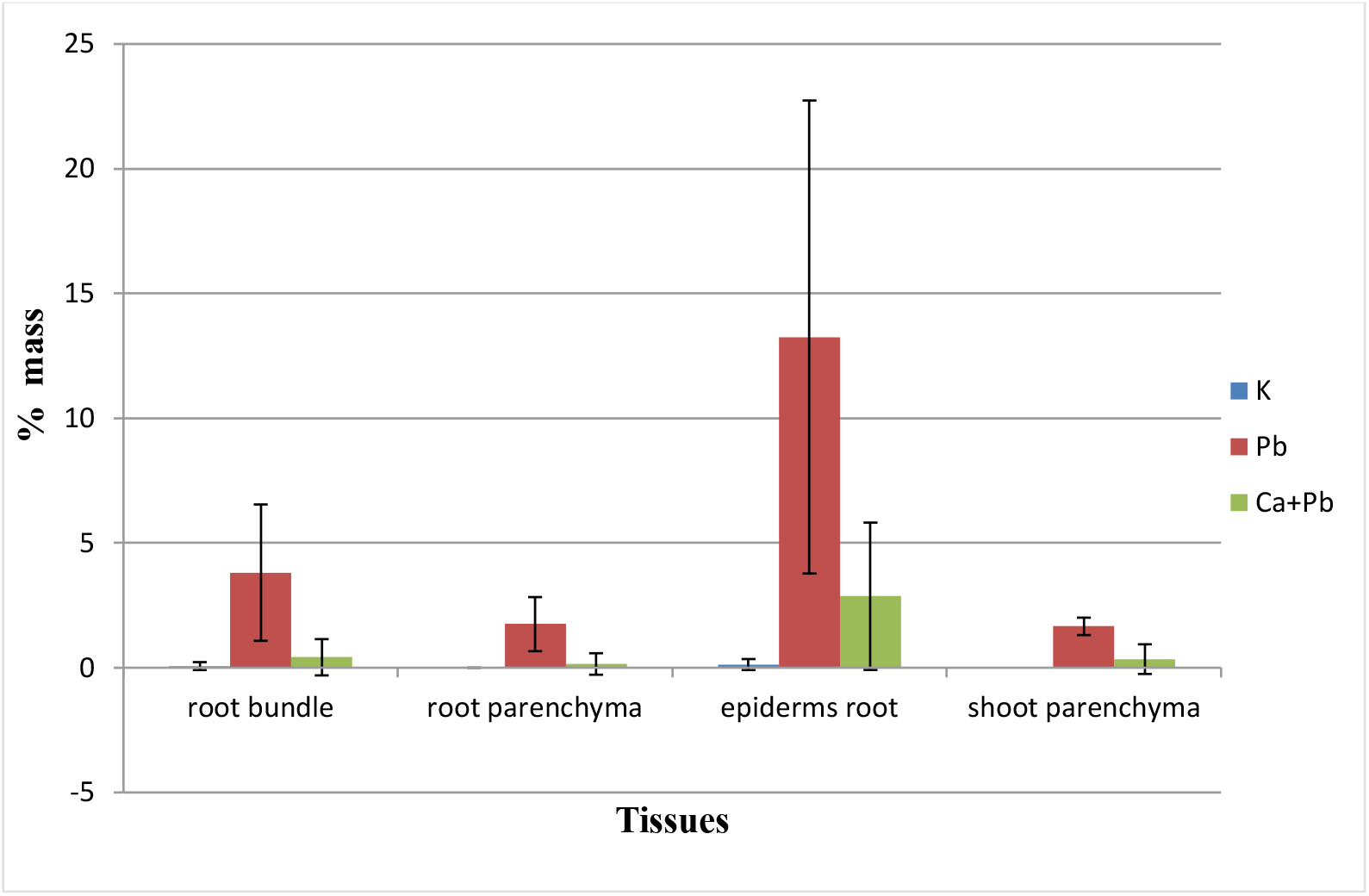
Lead concentrations in C. sativus L. tissues, roots and shoots treated with lead and calcium nitrate + lead. The figure is based on the numerical results from the regions analysed. No statistically significant differences.

### 3.8. Calcium foliar fertilisers

Since calcium nitrate reduces the uptake of lead by plants, it was also examined whether a commercially available calcium foliar fertiliser would have a similar effect. The results are shown in Fig. 14. It was found that in this case there was also a reduction in lead concentrations in *C. sativus* L. roots and shoots. After foliar application of 0.5% Viflo Cal S fertilizer (Agrosimex,2013), it was found that lead concentrations in the roots were decreased by 51% an in the shoots by 30% (Fig. 14). Finally, the amount of lead per plant was decreased on average from 65.6 µg to 30.9 µg.

**Fig. 14.**
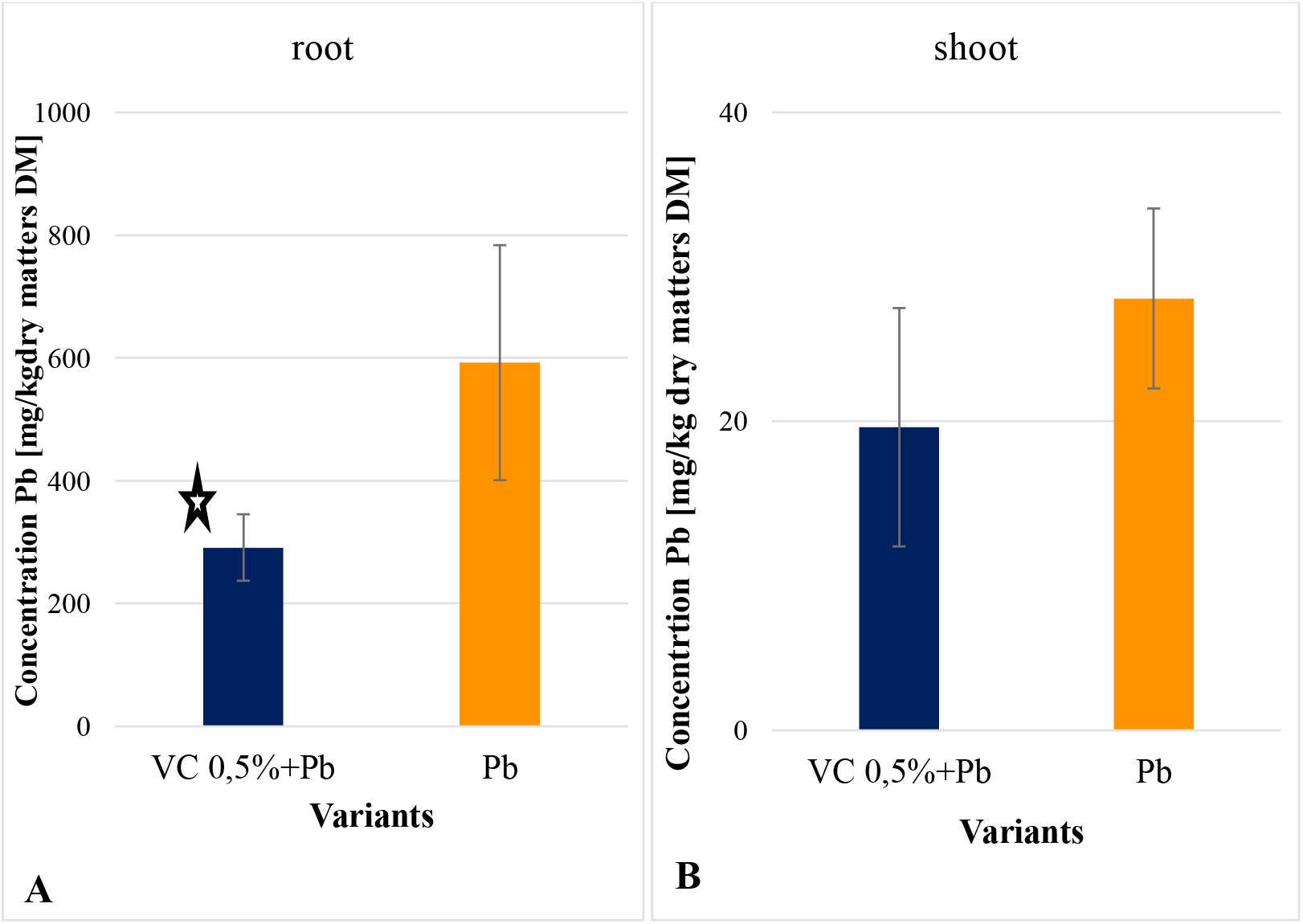
Lead concentrations in roots (A) and shoots (B) of *C. sativus* L. plants treated with lead administered to the roots and after foliar treatment of plants with Viflo Cal S fertiliser. Viflo Cal S preparation at 0.5%; lead at 3 mg/l. Asterisks indicate statistically significant differences with respect to lead. (Kruskal-Wallis test, significance level α = 0.05)

It was thus shown that the calcium fertiliser Viflo Cal S applied foliarly to *C. sativus* L. plants has the potential to reduce the uptake of lead by plants just like the InCa biostimulant and calcium nitrate. This finding indicates that calcium foliar fertilisers may have a similar effect, which requires further research.

## 4. DISCUSSION

As explained in the introduction, this paper is the first one that undertakes research leading to a reduction in lead uptake by crops using commercial biostimulant preparations. This study was conducted from baseline.

**4.1.** Firstly, it was necessary to select a suitable biostimulator for the study. Based on the *A. cepa* test, four commercial products were tested, which differed in terms of their composition (Table 1). The main difference was different hormonal composition (CaT, CalFlux) and different calcium concentration. Finally, it was found that all tested products reduced lead toxicity to epidermal cells of onion (*A. cepa* L.). In contrast, the InCa activator was most effective. That is why the InCa preparation was selected for further research.

**4.2.** The second step of this study examined which metals were susceptible to the InCa activator. The InCa activator was found to produce the strongest reductions in toxicity for lead (by 48%). The other divalent metal (Zn, Cd) by only about 20%. Based on this, lead was selected for all further studies. Hence, the proposed method can only be used to reduce lead concentrations in the plant.

**4.3.** In the next step, it was examined which crop species would be best suited for the study. The initial assumption of this study was that an increase in calcium concentrations in plants would result in a decrease in lead. This hypothesis finds support in the literature (as outlined in the Introduction).

In this situation, it was reasonable to build on the findings of Antosiewicz (1993), who found a relationship between the tolerance of various plant species to lead and their tolerance to calcium deficiency. In this way, four plant species were selected, differing in terms of their tolerance to lead and calcium deficiency. Finally, two species were selected for further research. The first species was tomato (*S. lycopersicum* L.), which had high tolerance to lead and good tolerance to calcium deficiency. In contrast, the second species was cucumber (*C. sativus* L.), which had medium tolerance to lead and medium tolerance to calcium deficiency.

In the authors’ opinion, the choice of these two species that differ in terms of their response to lead and calcium is a good representation of crops.

**4.4.** The next step was to test whether foliar application of the InCa activator to the plants would reduce lead uptake by plant roots. Concentrations of both InCa and lead were chosen so as not to cause interference with plant growth and development, bringing this study closer to natural conditions (Wierzbicka, 1995).

Finally, a reduction in lead uptake by plant roots was found to be most significant in tomato (*S. lycopersicum* L.), i.e. by 47%, and in cucumber (*C. sativus* L.) to 36%. An analysis of all the results obtained from the three plant species tested revealed that foliar application of InCa reduced lead concentrations in roots and in shoots by 44%.

The result obtained is definitely satisfactory, as it shows that it is possible to reduce the entry of lead into the biological cycle by approximately 44%.

The results of this study were confirmed by microscopic examination. The degree of root staining was observed after the use of histochemical method (dithizone). There was weak staining of the roots after treatment with InCa. This proves the reduced amount of lead that entered the plants, influenced by the foliar application of the InCa activator. On the other hand, at the cellular level, there was a reduction in the amount of lead in the root cell walls (apoplast) and a reduction in the amount of lead that entered the cytoplasm (symplast).

Therefore, it is possible that the InCa activator reduces lead transport via both symplastic and apoplastic pathways.

**4.5.** In an effort to go deeper into understanding the mechanism for the reduction of the uptake of lead by plants, it was examined which of its InCa activator components are responsible for this process. The InCa activator is a product with a complex chemical composition. It contains calcium nitrate, plant hormones, citric acid, etc.

Finally, calcium nitrate administered foliarly was found to reduce lead uptake by *C. sativus* L. cucumber plants by 72%. This result was confirmed by microscopic examination (scanning electron microscopy and X-ray microanalysis), which indicated a strong reduction in lead concentrations in all tissues, plant shoots – especially in the vascular bundles – and in the cells of the cortex. A similar result was obtained after the application of the commercial calcium foliar fertiliser Viflo CalS, which reduced the amount of lead in the roots *C. sativus* L. by approximately 51%.

Based on the above-mentioned studies, it was assumed that calcium nitrate included in the InCa activator mainly causes a reduction in the uptake of lead by plants. The commercial foliar fertiliser revealed a similar effect, so it is very likely that other calcium fertilisers will have a similar action. Research in this direction will continue.

This study indicates that there is potential to reduce lead uptake by crops by approximately 44%. The proposed method is environmentally beneficial as the products applied foliarly are in low concentrations. Calcium transport activators (e.g. InCa) and calcium nitrate have the potential for harmless effect. So far, no method was developed to reduce lead concentrations in crops. Therefore, the result obtained in this study is very satisfactory. This is cutting-edge research that was decribed for the first time in this paper.

## 5. Conclusions

For the first time, research was attempted to protect plants from lead ingress, which has direct implications for the amount of lead in the biological cycle and in our diet.

Studies revealed that foliar application of agricultural biostimulants that are calcium transport activators (e.g. InCa, Plant Impact company) can reduce lead concentrations in plants by approximately 44%.

Such beneficial effects of the calcium transport activator InCa were found to be caused by the calcium compounds included in this preparation. A similar effect was obtained after foliar treatment of the plants with calcium nitrate and with a commercially available calcium fertiliser.

Hence, for the first time, the existence of a possibility to reduce lead uptake by plants was indicated. According to the authors, a 44% effect is considered very favourable. The indicated pathway can be used for further research and, in the long term, for application studies. In doing so, it should be taken into account that the possibility of reducing the uptake of lead by the plant may apply from the first stages of plant growth and development, i.e. from the moment when the plants are exposed to lead uptake from the soil. In contrast, this method will not be applicable when lead has already been taken.

## Acknowledgements

The research was funded by NCN grant no. 2016/21/B/NZ8/01564

